# How fast are cells dividing: Probabilistic model of continuous labeling assays

**DOI:** 10.1101/550574

**Authors:** Julian Rode, Torsten Goerke, Lutz Brusch, Fabian Rost

## Abstract

Correct estimates of cell proliferation rates are crucial for quantitative models of the development, maintenance and regeneration of tissues. Continuous labeling assays are used to infer proliferation rates *in vivo*. So far, the experimental and theoretical study of continuous labeling assays focused on the dynamics of the mean labeling-fraction but neglected stochastic effects. To study the dynamics of the labeling-fraction in detail and fully exploit the information hidden in fluctuations, we developed a probabilistic model of continuous labeling assays which incorporates biological variability at different levels, between cells within a tissue sample but also between multiple tissue samples. Using stochastic simulations, we find systematic shifts of the mean-labeling fraction due to variability in cell cycle lengths. Using simulated data as ground truth, we show that current inference methods can give biased proliferation rate estimates with an error of up to 40 %. We derive the analytical solution for the Likelihood of our probabilistic model. We use this solution to infer unbiased proliferation rate estimates in a parameter recovery study. Furthermore, we show that the biological variability on different levels can be disentangled from the fluctuations in the labeling data. We implemented our model and the unbiased parameter estimation method as an open source Python tool and provide an easy to use web service for cell cycle length estimation from continuous labeling assays (https://imc.zih.tu-dresden.de/cellcycle).

## Introduction

Cells arise from cells by cell division. The timing of cell divisions has to be tightly regulated to ensure proper development and maintenance of tissues. Consequently, precise estimates of cell cycle lengths are necessary to understand tissue development and maintenance, e.g. to access the contribution of cell divisions to the growth of developing and regenerating tissues (Chara et al., 2014; Rost et al., 2016; Jülicher & Eaton, 2017) or to access how environmental changes influence neurogenesis (Overall et al., 2016).

A widely used method for inferring cell cycle lengths *in vivo* is the continuous labeling assay (Nowakowski et al., 1989; Blanpain & Simons, 2013). In those assay, cells — even deep inside an adult body not accessible by microscopy techniques — are continuously exposed to a label that is incorporated into newly synthesized DNA (Figure 1). Intuitively, one expects that a faster cell cycle leads to a faster uptake of the label in the cell population. A quantitative mathematical model of the labeling assay allows to estimate the proliferation rate from the measurement of the number of labeled cells at successive time points.

**Figure 1:**
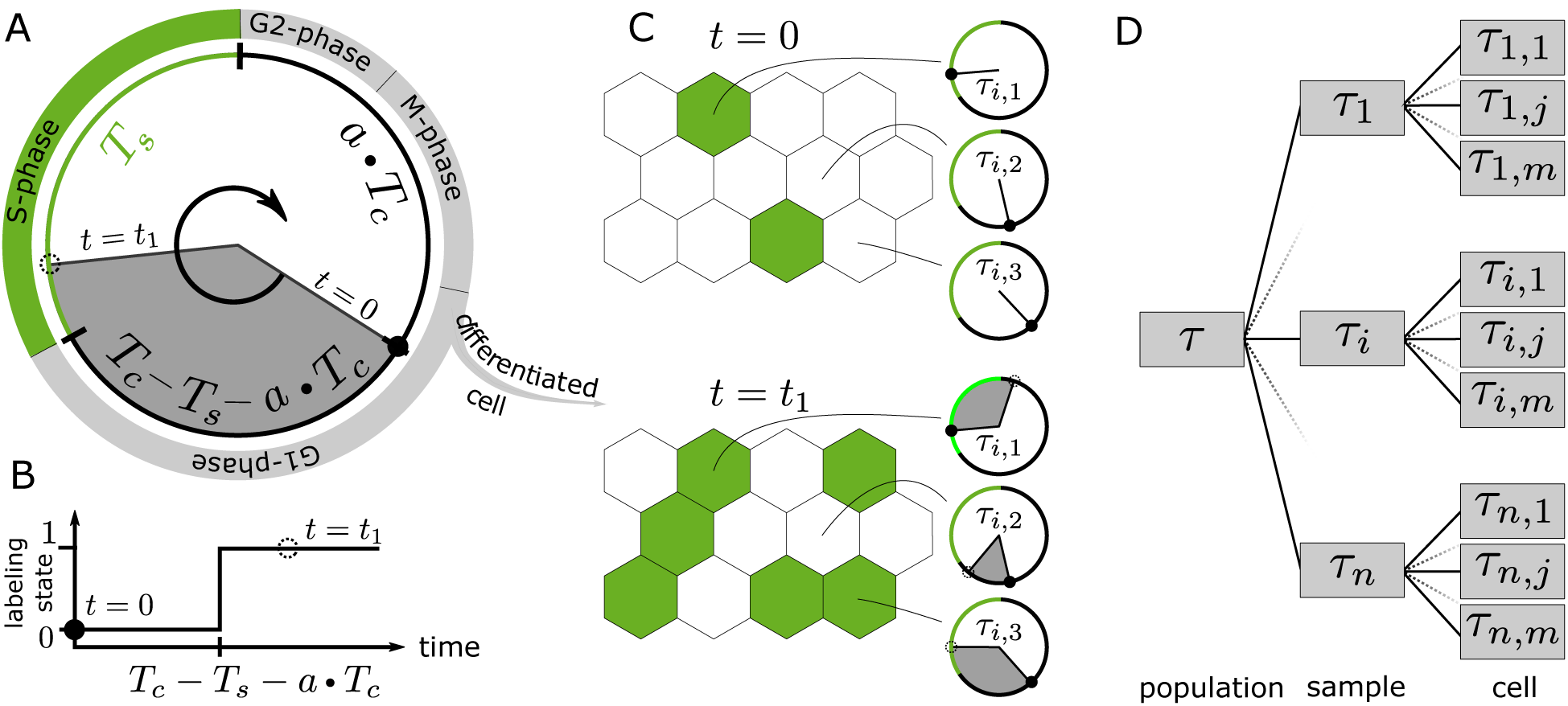
Model for continuous labeling assays. **A**: The outer and inner ring visualize the biological cell cycle and our single cell model. During the S-phase, shown in green, DNA is synthesized and incorporates the label. In contrast to the biological cell age, the model cell age *aT_c_* starts at the end of the S-phase. The gray shaded area indicates the time the cell is exposed to the labeling analog which is *T_c_− T_s_−aT_c_*. When this gray interval reaches into the S-phase, that cell is labeled. B: Time evolution of the single cell model. The labeling state changes from zero (unlabeled) to one (labeled) at *t* = *T_c_− T_s_− aT_c_*. C: Cartoon of the labeling assay for tissue shows labeled cells (green) at the start of the experiment (*t* = 0) and later (*t* = *t*_1_). For three exemplary cells, their individual cell cycle is shown. Since they have different cell ages and cell cycle lengths, they become labeled at different times. Note, cell *j* = 2 is older than cell *j* = 3, but stays unlabeled due to its significantly longer cell cycle length *τ_i_*,_2_ which is represented by the smaller gray area compared to the other two cells. D: Hierarchical probabilistic model for the cell cycle length. Each cell has an individual cell cycle length *τ_i,j_* which depends on the sample level cell cycle length *τ_i_*, which in turn depends on the population level cell cycle length such that *< τ_i,j_ >_j_* = *τ_i_*, *< τ_i,j_ >_i,j_* =*< τ_i_ >_i_*= *τ*.

The majority of previously developed models for labeling assays are deterministic (Nowakowski et al., 1989; Bernard et al., 2010; Lefevre et al., 2013; Schittler et al., 2013; Hross & Hasenauer, 2016). These models fall in to two classes: While some model the state of cells with a continuous variable for label concentration, others model the state just with a binary variable, i.e. a cell is either labeled or not labeled (Nowakowski et al., 1989; Lefevre et al., 2013). The latter approach is more suitable for *in vivo* systems as the precise amount of label that is administered and the distribution of the label in the body are uncertain. Therefore, we here consider the case of binary cell states. For this binary case, the widely used model by Nowakowski et al. (1989) assumes asymmetric, asynchronous cell divisions, Lefevre et al. (2013) extended this model towards symmetric, possibly synchronized divisions.

We were concerned about several assumptions that are used in those deterministic models. First, the models by Nowakowski et al. (1989) and Lefevre et al. (2013) assume that all considered cells have identical cell cycle phase lengths. However, cell cycle phase lengths vary between cells as shown by time-lapse lineage tracing experiments (Bach et al., 2014). Second, the models assume identical average cell cycle lengths of different tissue samples. Continuous labeling assays cannot provide actual time-series data, because samples have to be fixed for measurements. It seems reasonable to assume that the average proliferation rate also varies between different samples. Third, an additive Gaussian error model for the number of labeled cells is assumed to perform parameter estimation. While this assumption seems reasonable, it adds the measurement errors as free parameters in the estimation procedure. Importantly, this assumption breaks when nearly no or nearly all cells are labeled.

It is thus of interest to learn how a probabilistic model of labeling assays would compare to the deterministic model predictions. In related fields, probabilistic models have previously shed light on essential biological processes that remained poorly understood based on deterministic approaches (Elowitz et al., 2002; McDonnell & Abbott, 2009; Robb & Shahrezaei, 2014). It is important to understand how biological variability and the measurement error lead to fluctuations in the measured number of labeled cells (Oates et al., 2009). We hypothesized that neglecting this variability in the model could lead to biased estimates of cell cycle lengths. Furthermore, we asked whether the explicit modeling of the variability and measurement error allows estimating the cell cycle length more accurately and precisely. Further, we wondered whether the biological variability on cell and tissue scale could be estimated and disentangled given the fluctuating data in the labeling assays.

Here, we develop a probabilistic model of continuum labeling assays that incorporates biological variability on the cell and sample levels as well as the measurement errors. By simulating this model, we find systematic deviations from the previous deterministic model in the fraction of labeled cells. To access whether these deviations are of practical relevance for the estimation of the cell cycle length, we derive an analytical expression for the likelihood. Using this expression, we perform a parameter recovery study based on the maximum likelihood method. We compare the accuracy and precision of our model to the previous deterministic model. Finally, we present a web service that allows to easily analyze continuous labeling assays with our probabilistic model.

### Probabilistic Model of S-Phase Labeling

In order to develop a probabilistic model accounting for variability between cells (indexed by *j*) and between samples (indexed by *i*), we first define a single cell model of S-phase labeling. As we consider asymmetric cell divisions the total number of cycling cells stays constant and the actual division of a cell after M-phase is not relevant. Instead label incorporation during the S-phase is important. Therefore, we consider the end of the previous S-phase as our start time for the cell age. When the cell enters its S-phase, the label, e.g. a Thymidine analog like BrdU or EdU, becomes embedded in the newly synthesized DNA and the cell gets labeled. Then, the labeling state *s_i,j_* of cell *j* in sample *i* is a Heaviside function *𝓗* which changes the state from zero (unlabeled) to one (labeled):

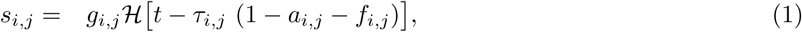

where *g_i,j_* ∈ {0, 1} determines if that cell proliferates, *τ_i,j_* is the cell cycle time of that cell (*T_c_*), *f_i,j_* ∈ [0, 1] the relative S-phase length of that cell (*T_s_/T_c_*), *a_i,j_* ∈ [0, 1] the age of that cell at *t* = 0 normalized to *T_c_* and *t* the time after the start of the labeling experiment (Figure 1A,B,C).

In the idealized limit of identical cellular parameters and uniformly distributed cell ages *a_i,j_*, we recover the Nowakowski model:

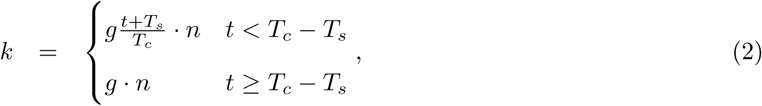

where *k* is the expected number of labeled cells in a sample of *n* cells and *g* the probability of a cell to be proliferating. In this limit, the fraction of labeled cells *k/n* in our model equals the labeling index of the Nowakowski model.

In the biological more relevant probabilistic case, we define random variables for the other cell parameters: *τ_i,j_* and *g_i,j_*. Since we consider the phase stretching model (Dowling et al., 2014) of the cell cycle *f_i,j_* = *f* is independent of individual cells and thus not a random variable. The Boolean parameter *g_i,j_* is determined by a Bernoulli distribution *𝓑𝓔*(*g*) with the tissue-scale growth fraction *g* as success probability. For the cell cycle times *τ_i,j_*, we assume that cells in each sample are more similar than cells from different samples and that the samples are independent. Therefore we use a hierarchical probabilistic model with two levels (Figure 1D): the sample level (first random variable) and the cell level (second random variable dependent on the first). Hence, the random variables for the cell level are not independent and identically distributed, but depend on the realization of the sample level random variable. We choose a log-normal distribution for the two levels, since it corresponds well to experimental data showing an asymmetric cell cycle length distribution with a long tail (Moussy et al., 2017). Thus we have the sample level random variable *T* and a set of random variables *T_i_* for the cell level in sample *i*. Using these random variables, the single cell model (Eq. 1) can be extended to a probabilistic model:

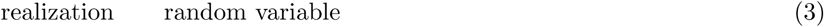

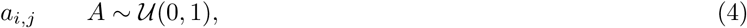

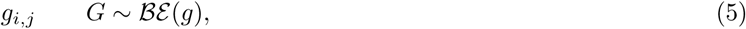

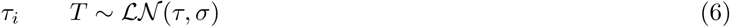

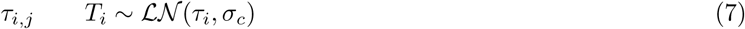

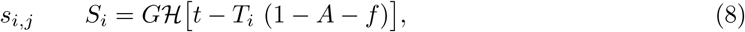

where *S_i_* is the random variable for the labeling state. Since *G* and the Heaviside function are Boolean, *S_i_* is also Boolean and thus it follows a Bernoulli distribution.

## Results

### Model Simulation

For each sample we draw stochastic cell parameters from the random variables: *T_i_*, *G* and *A* and 10000 samples per measurement time. We scale the time to the true cell cycle time which will be set to one without loss of generality. Then the histogram, percentiles and mean of labeled cells can be calculated for each time point. The numerical solutions for the probabilistic model is shown in Figures 2 and 3. The width of the distribution of the labeling index starts narrow at *t* = 0, increases to its maximum around *t* = 0.5 and then decreases in the plateau. Importantly, its mean values agree with the Nowakowski solution only at the start and at the end. For intermediate times, Nowakowski’s solution is first lager and later smaller than the true data. Whether this discrepancy is relevant is checked with an exemplary parameter recovery study using the 5 time points marked with black triangles in Figure 2. The recovered parameters turned out to differ from the ground truth values of the simulation (Figure 2) which demonstrates that the Nowakowski model applied to realistic data can yield a significant bias.

**Figure 2:**
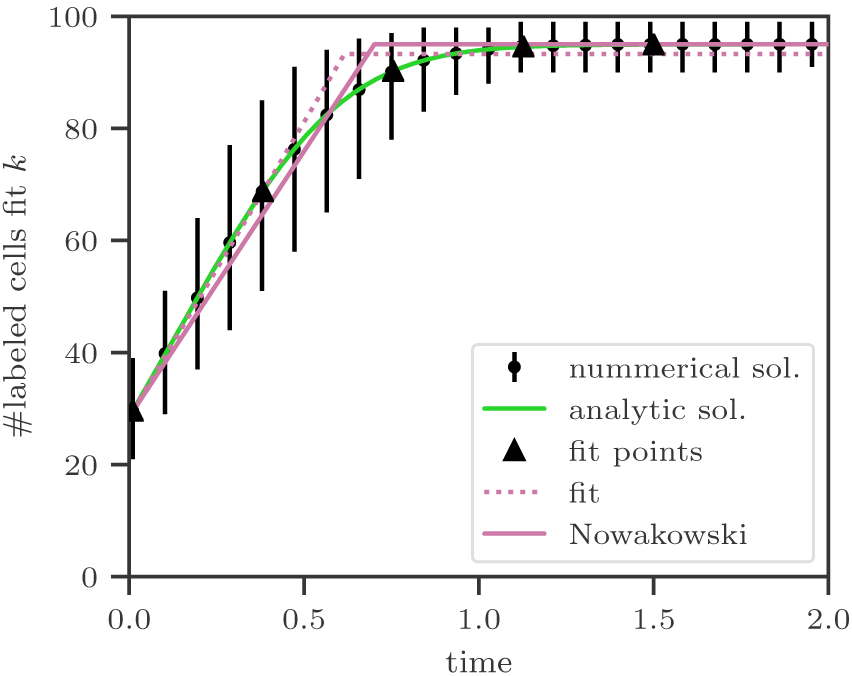
Labeling time courses. For the probabilistic model, the mean number of labeled cell is plotted with black dots and the error bars indicate the 95 percentile region. The green and red solid curves show the analytic solutions for identical parameter values of our probabilistic model (Eq. 15) and Nowakowski’s model (Eq. 2). The dotted red line shows the fitted Nowakowski model using the black triangles as data, resulting in the point estimates: *τ* = 0.87 *±* 0.05, *f* = 0.31 *±* 0.02 and *g* = 0.93 *±* 0.01. Only the true value for the S-phase fraction lies inside its error interval, but for the other two parameters their error intervals do not include the true value. For the simulation we used 10000 samples per time point and *n_i_* = 100 cells per sample. The true parameter values are: *τ* = 1, *f* = 0.3, *g* = 0.95, *σ_c_*= 0.3 and *σ* = 0.2.

**Figure 3:**
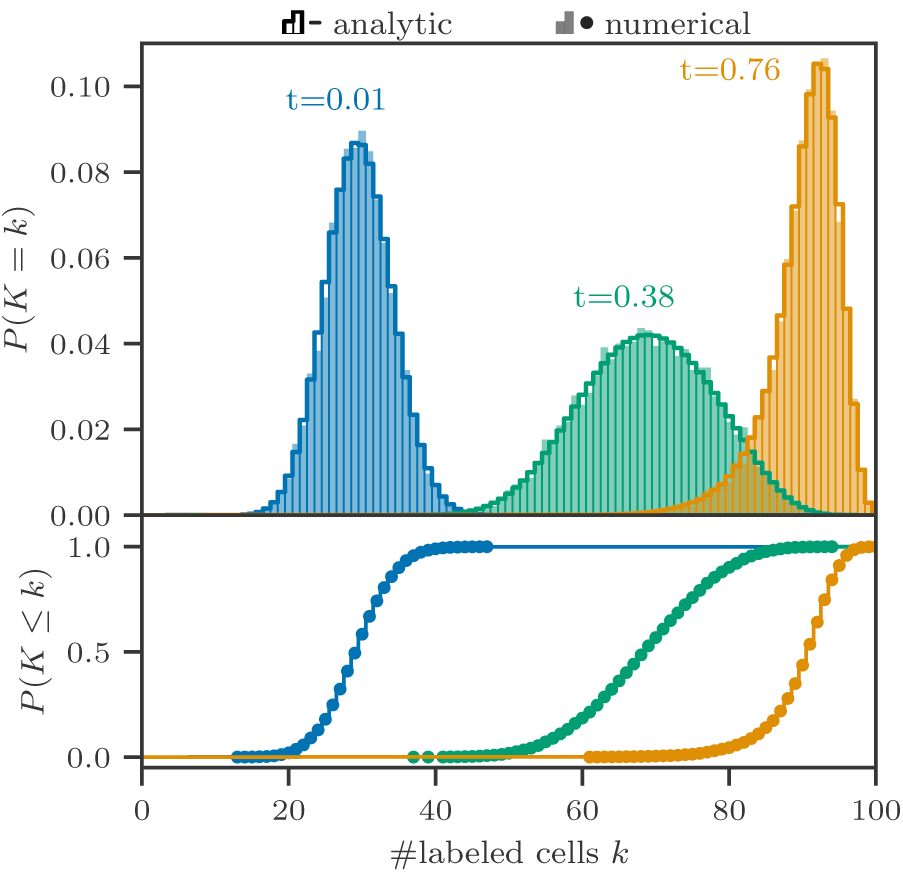
Comparison of analytical solution to the simulation for histograms and cumulative distribution functions. A: For 3 time points our analytic solution of the probabilistic cell labeling model (solid curve) is compared to the histograms of the numerical solution (shaded area). B: The corresponding cumulative distribution functions are shown. For parameters see Figure 2.

### Analytic Expression for the Likelihood

In order to calculate the likelihood for our probabilistic model we need to determine the probability mass function (pmf) of the labeling state for one sample *S_i_* and generalize it to the pmf for an arbitrary sample *S*. Since the Boolean labeling state *S_i_* must follow a Bernoulli distribution, the expectation value of *S_i_* equals its probability *p*(*τ_i_, g, f, σ_c_*; *t_i_*). Therefore, the probability for a cell to be labeled in the sample *i* is given by:

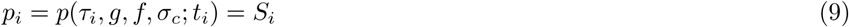

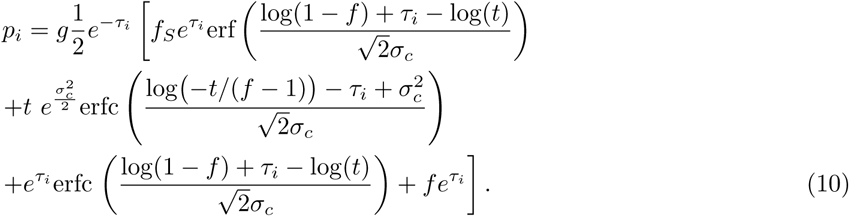

In the probabilistic case, the counting process of labeled cells itself needs to be considered because the number of observed cells is relatively small which introduces noise. Here, it is modeled by a Binomial process *𝓑*(*n_i_, p_i_*). The number of labeled cell in the sample *i* equals the number of successes in a Binomial process using *n_i_* trials and *p_i_* as success probability, where *n_i_* corresponds to the number of cells observed in one sample:

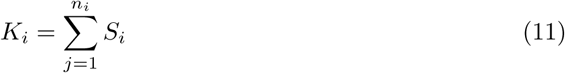

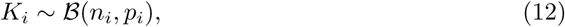

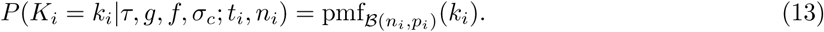

In order to generalize this distribution for different samples, it is interpreted as the conditional probability *P* (*K* = *k|T* = *τ_i_*) of finding one sample *i* in the overall experiments. Consequently, the law of total probability can be applied in order to calculate the probability to find *k* labeled cells independent of a specific sample:

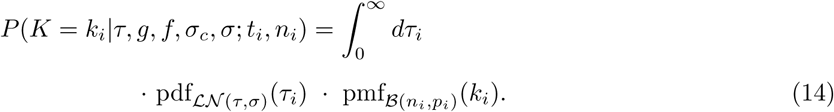

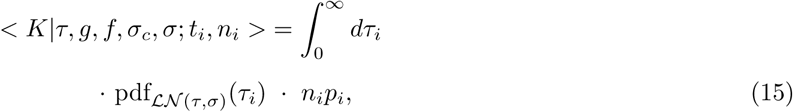

Figure 3 depicts the perfect match between the probabilistic simulation and the analytic solution (Eq. 14) of our probabilistic model. The pmf solves the inverse problem and depends on the following set of parameter: *τ, g, f, σ_c_, σ*. The pmf with the parameters *ϕ* and the experimental data **k** = *{k*_0_*, k*_1_*, …, k_n_}* at **t** = *{t*_0_*, t*_1_*, …, t_n_}*) build the likelihood function *ℒ* for the cell labeling experiment:

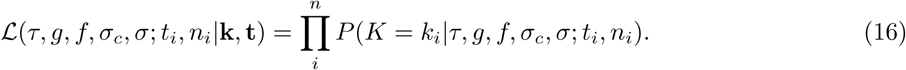

### Parameter Recovery Study

To obtain parameter estimates with the likelihood (Eq. 16) and test its accuracy, we preform a parameter recovery study under realistic conditions. We simulate an experiment as in Figure 2, but instead of 10000 samples per time point we consider two cases with 5 and 100 samples per time point. Our likelihood and a least-square fit for the Nowakowski model are used in parallel to fit these simulated experiments. We use the software package ‘minuit’ to minimize the likelihood and calculate the 95% error intervals using the build-in parabolic error estimate. Thereby, the Hessian matrix is numerically calculated and its entries are used to fit a parabolic error profile around at the point estimate. Using 1000 repetitions of the point estimation, the distributions of the parameter estimates are determined and their bias as well as their accuracy are calculated. In Figure 4 the resulting histograms for the point estimates are depicted for both cases. We again encounter the bias of the Nowakowski model (red) which does not vanish in the high sample number case despite a narrower distribution. In contrast, our likelihood has no bias and in the high sample case increases the accuracy as one would expect. Additionally, in the high sample number case it is possible to infer the sample and cell level variability *σ* and *σ_c_* very well. In both cases the accuracy is close to the expected 95% using our probabilistic model opposed to the underestimated error in the Nowakowski model (Table 1).

**Table 1:**
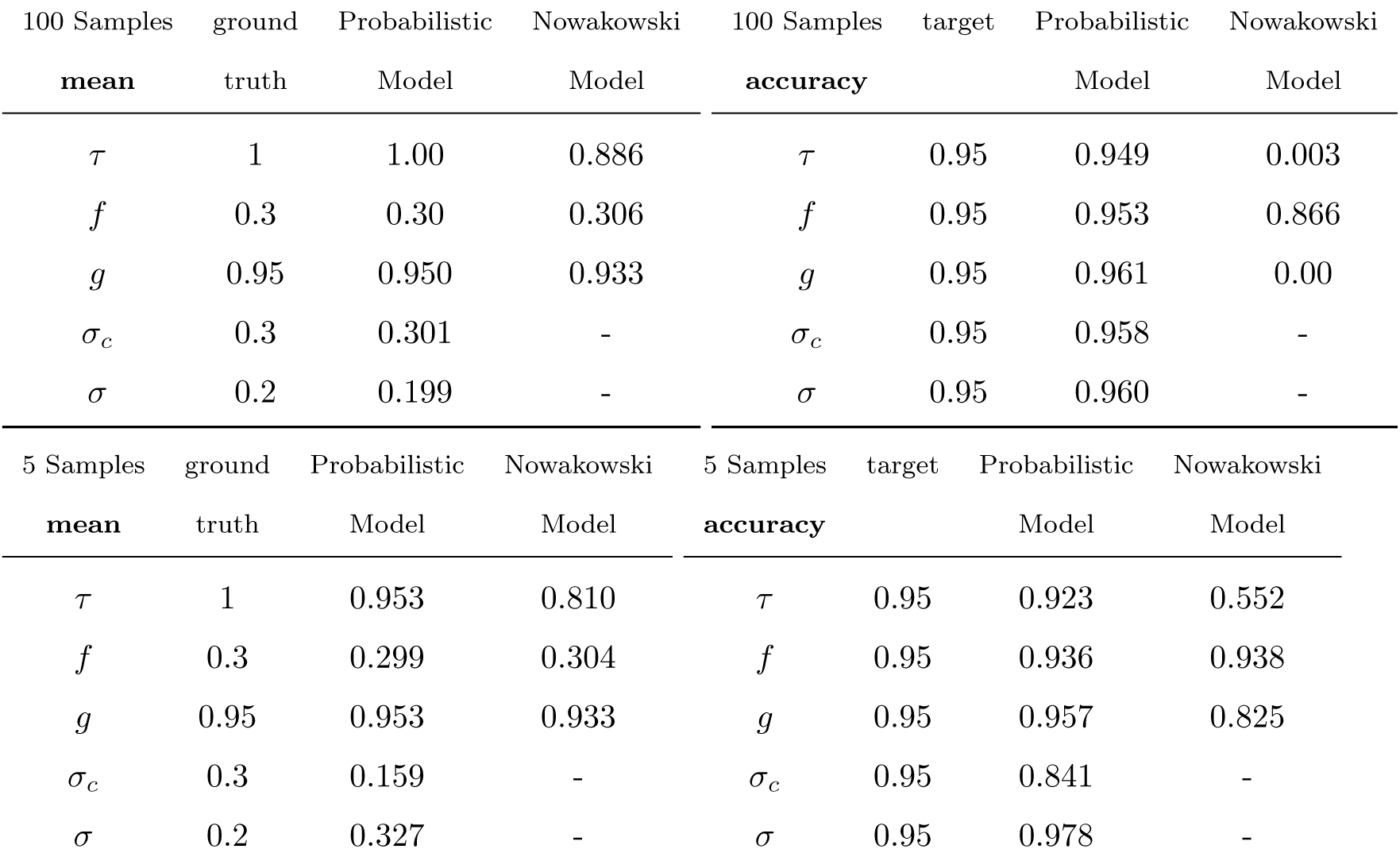
Summery of the results of the parameter recovery study. For the cases with 5 and 100 samples per time point the mean point estimate and the accuracy using 1000 trials are presented.

| 100 Samples | ground | Probabilistic | Nowakowski | 100 Samples | target | Probabilistic | Nowakowski |
| --- | --- | --- | --- | --- | --- | --- | --- |
| **mean** | truth | Model | Model | **accuracy** |  | Model | Model |
| $\tau$ | 1 | 1.00 | 0.886 | $\tau$ | 0.95 | 0.949 | 0.003 |
| $f$ | 0.3 | 0.30 | 0.306 | $f$ | 0.95 | 0.953 | 0.866 |
| $g$ | 0.95 | 0.950 | 0.933 | $g$ | 0.95 | 0.961 | 0.00 |
| $\sigma_c$ | 0.3 | 0.301 | - | $\sigma_c$ | 0.95 | 0.958 | - |
| $\sigma$ | 0.2 | 0.199 | - | $\sigma$ | 0.95 | 0.960 | - |

| 5 Samples | ground | Probabilistic | Nowakowski | 5 Samples | target | Probabilistic | Nowakowski |
| --- | --- | --- | --- | --- | --- | --- | --- |
| **mean** | truth | Model | Model | **accuracy** |  | Model | Model |
| $\tau$ | 1 | 0.953 | 0.810 | $\tau$ | 0.95 | 0.923 | 0.552 |
| $f$ | 0.3 | 0.299 | 0.304 | $f$ | 0.95 | 0.936 | 0.938 |
| $g$ | 0.95 | 0.953 | 0.933 | $g$ | 0.95 | 0.957 | 0.825 |
| $\sigma_c$ | 0.3 | 0.159 | - | $\sigma_c$ | 0.95 | 0.841 | - |
| $\sigma$ | 0.2 | 0.327 | - | $\sigma$ | 0.95 | 0.978 | - |

**Table 2:** Symbol definitions

| Symbol | Definition |
| --- | --- |
| $T_c$ | Cell cycle time |
| $T_s$ | S phase time |
| $a$ | cell age |
| $g$ | growth fraction |
| $i$ | samples index |
| $j$ | cell index |
| $n_i$ | number of cells in a sample $i$ |
| $n$ | total number of samples |
| $t$ | time since labeling started |
| $\tau_{i,j}$ | cell cycle time of cell $i$ from sample $j$ |
| $\tau_i$ | mean cell cycle time of sample $j$ |
| $g_{i,j}$ | proliferation state of cell $i$ from sample $j$ |
| $s_{i,j}$ | labeling state of a cell $i$ from sample $j$ |
| $p_i$ | probability for a cell in sample $i$ to be labeled |
| $k_i$ | number of labeled cells in samples $i$ |
| $a_{i,j}$ | normalized age of the cell at $t = 0$ |
| $f_{i,j} = f$ | S-phase fraction |
| $T_i$ | random variable for mean cell cycle length of sample $i$ |
| $T$ | random variable for cell cycle length |
| $S_i$ | random variable for labeling state of cells $i$ from sample $j$ |
| $A$ | random variable for cell age |
| $G$ | random variable for proliferation state |
| $K$ | random variable for number of labeled cells |
| $\mathcal{BE}(p)$ | Bernoulli distribution with probability $p$ |
| $\mathcal{B}(n, p)$ | Binomial distribution with $n$ trials and probability $p$ |
| $\text{pmf}_{\mathcal{B}}(k)$ | Probability mass function of the Binomial distribution |
| $\mathcal{U}(a, b)$ | Uniform distribution from $a$ to $b$ |
| $\mathcal{LN}(\mu, \sigma)$ | log normal distribution with mean $\mu$ and variance $\sigma$ |
| $\text{pdf}_{\mathcal{LN}}(\tau)$ | Probability density function of the log normal distribution |
| $\sigma_i^2 = \sigma_c^2$ | cell to cell variance |
| $\sigma^2$ | sample to sample variance |

**Figure 4:**
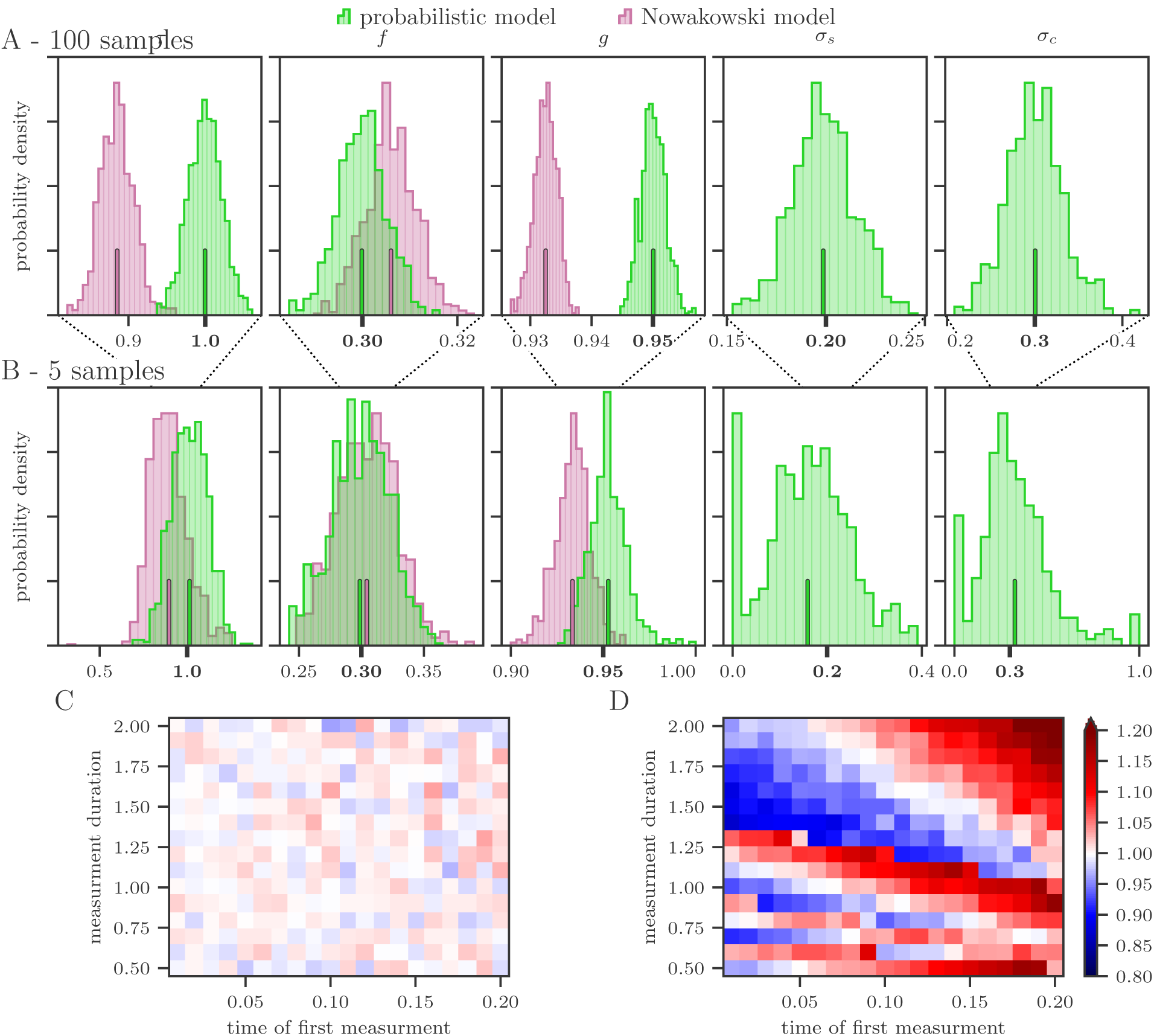
Parameter recovery study. The green histograms show the point estimates using our likelihood (Eq. 16) and the red ones show the point estimated from a least-square fit using the Nowakowski model. The correspondingly colored large tick indicate the mean value and the ground truth is marked by the bold ticks. A: 100 samples per time point B: 5 samples per time point. C and D: For different equidistant placements of the five measurement times, the point estimates of the cell cycle *T_c_*are shown for our probabilistic model C and the Nowakowski model D. The parameters are as in Fig 2.

The placement of the five time points could have an influence on the point estimates. To evaluate the influence of the choice of measurement times, we analyze the distance and initial offset of five equally spaced time points using 10 repetitions. In fig. 4C,D, results of this evaluation are presented which show the averaged estimated cell cycle length *τ*. For the used interval of measurements points, our probabilistic model yields equally good results independent of the measurement points. The cell cycle varies slightly around the true value with a standard deviation of 0.04. In contrast, the bias of Nowakowski’s model can change drastically (up to 40%) even for a small variation of the measurement points. It ranges from a maximum cell cycle length of 1.25 to a minimum of 0.87.

The biologically founded variability of the cell cycle on cell and sample level systematically changes the averaged number of labeled cells leading to a bias of the point estimates using Nowakowski’s deterministic model. Since the labeling assays are extensive experiments there is usually a small number of measurements which increases the imprecision of the point estimates. Therefore, the spread of the point estimates can be larger than the bias. But with our probabilistic model solution an increase in the numbers of measurements at the given time points will increase the precision of the point estimate in contrast to Nowakowski’s model where just the bias will appear more clearly. Additionally, our model is able to produce accurate error intervals resulting in the targeted accuracy independent of the number of measurements (Table 1) and these characteristics only change very slightly with the placement of the measurement points.

### Web service for parameter estimation

Since the parameter estimation with our probabilistic model and its likelihood (Eq. 16) is not as straight-forward as with the Nowakowski model, we developed an easy to use web service (https://imc.zih.tu-dresden.de/cellcycle) for the parameter estimation. The user can simply insert the experimental data manually or upload a CSV file containing the data (Figure 5). All user data are only temporally stored on our server as long as they are needed for the parameter estimation.

**Figure 5:**
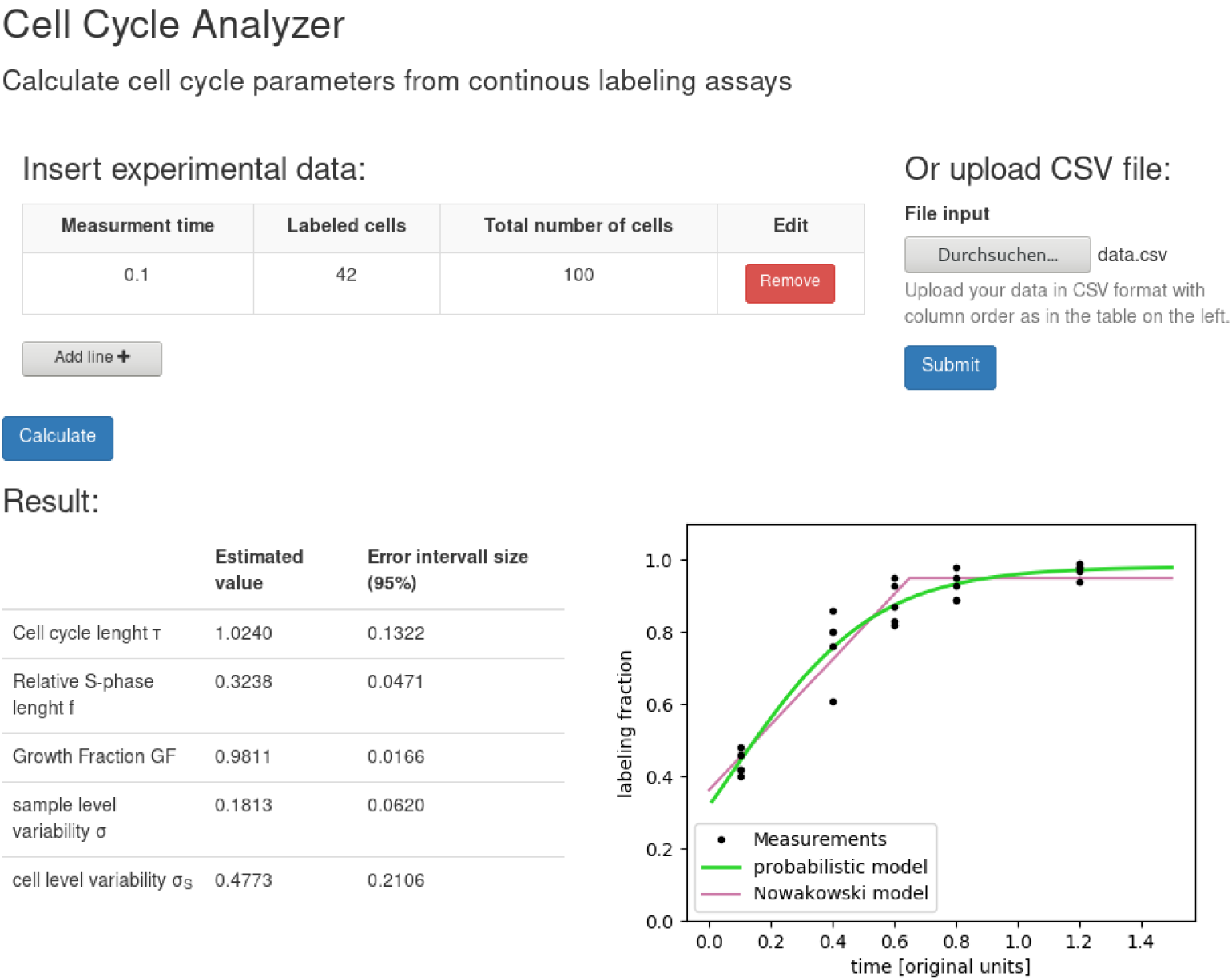
Screenshot of the web service. The screenshot illustrates our web service (https://imc.zih.tu-dresden.de/cellcycle) for parameter estimation. In the example, (simulated) data are uploaded via the CSV file upload, but data can also be entered manually. The interfered parameters are displayed along with a plot of the normalized data, our probabilistic model and the Nowakowski model.

The resulting point estimates and the size of the 95% error interval are presented in a table as well as visualized in the figure. The figure shows a plot of the normalized data, our probabilistic model and the Nowakowski model. For our probabilistic model, the presented point estimates are applied to calculate the expected labeling fraction which is plotted in green. A separate fit for the Nowakowski model is preformed and used plotted in red.

Since the model is invariant under any time scale, the user just has to enter all times in the same units which equals the one of the output parameters.

## Discussion

We developed a probabilistic model of continuous labeling assays that takes into account biological variability of cell cycle lengths at the cell and sample level. Using Monte-Carlo simulations, we showed that this variability leads to systematic deviations in the prediction of the average number of labeled cells. We gave an exact analytical expression for the likelihood in the asymmetric division case which we used to perform a parameter recovery study. The parameter recovery study confirmed that the use of the deterministic model by Nowakowski can lead to a bias in cell cycle length estimation of up to 40%. In contrast, with our probabilistic model, we obtained unbiased estimates and correct credibility levels. Furthermore, our model allows to estimate and discriminate the contributions of variability on cell and sample level.

When estimating parameters in practice, the simplicity of the model by Nowakowski is appealing as it is much easier to implement compared to our likelihood (Eq. 16). We provide a web interface to the implementation of our probabilistic model at https://imc.zih.tu-dresden.de/cellcycle that should make parameter estimation easy.

In the limit of identical cell cycle phase lengths for all cells, our model prediction for the mean number of labeled cells converges to the deterministic prediction of Nowakowski et al. (1989). This verifies the consistency with the established model. The only other probabilistic modeling approach of the asymmetric, asynchronous case was presented by Blanpain & Simons (2013), which is based on a simple Markov birth process. This model does not address sample-level variability. Furthermore, it is not clear *a priori* whether using a Markov process for cell divisions is justified given that the length of continuous labeling assays is in the order of the cell cycle length itself (Klein et al., 2007).

For our model, we used a number of assumptions that should be considered when applying it to experimental data. First, we use the cell phase stretching model (Dowling et al., 2014). Other modes of cell phase length variability have been reported (Smith & Martin, 1973; Castor, 1980; Wellard et al., 2011; Yates et al., 2017) and their influence on cell cycle length estimates remains to be elucidated. Second, we decided to model cell cycle length distributions with a log-normal distribution. Other distributions, like the gamma distribution or normal distribution have been suggested (Scherf et al., 2012; Míguez, 2015). Furthermore, potential sub-populations of considered cycling cells will lead to multimodal parameter distributions. Again, the effect of our specific choice remains to be elucidated.

We derived an analytical expression for the likelihood (Eq. 16) only for the case pf asymmetric cell division. Considering asymmetric cell division is well justified for homeostatic stem cell populations. For expanding cell populations one should consider symmetric, self-renewing cell divisions (Rost et al., 2016). While the extension of our probabilistic towards symmetric cell divisions is straightforward, it is unclear whether one can find a tractable likelihood in this case. Although we did not give a tractable likelihood for this case, approximate Bayesian computation, probabilistic programming or other likelihood-free methods could be used for statistical inference in this case (Didelot et al., 2011; Klinger et al., 2018).

Our probabilistic model of continuous labeling assays allows to estimate precisely and accurately average cell cycle lengths as well as cell cycle length variability *in vivo*. This also exemplifies, how probabilistic models in general allow to estimate variability on cell and sample levels from statistical ensemble measurements. From a mathematical perspective, this work demonstrates how to disentangle individual stochastic contributions from cumulative ensemble data.

## Acknowledgments

We thank Anja Voß-Böhme, Osvaldo Chara, Jörn Starruß and Steffen Rulands for stimulating discussions. We acknowledge generous allocation of computer time by the Center for Information Services and High Performance Computing (ZIH) at TU Dresden.

## Author contributions

F.R. conceived the project. J.R. and F.R. developed and implemented the model and derived the analytical results. J.R. performed model simulations, data analysis and generated figures. T.G. developed the web service. J.R., T.G., L.B. and F.R. interpreted the data and wrote the manuscript. All authors gave their final approval for publication.

## Data and materials availability

All code used in this work is available in Python as Open Source from https://github.com/fabianrost84/clapy.

